# Size-dependent eco-evolutionary feedbacks in fisheries

**DOI:** 10.1101/2020.04.03.022905

**Authors:** Edeline Eric, Loeuille Nicolas

## Abstract

Harvesting may drive body downsizing along with population declines and decreased harvesting yields. These changes are commonly construed as consequences of direct harvest selection, where small-bodied, early-reproducing individuals are immediately favoured. However, together with directly selecting against a large body size, harvesting and body downsizing alter many ecological features, such as competitive and trophic interactions, and thus also indirectly reshape natural selection acting back on body sizes through eco-evolutionary feedback loops (EEFLs). We sketch plausible scenarios of simple EEFLs in which one-dimensional, density-dependent natural selection acts either antagonistically or synergistically with direct harvest selection on body size. Antagonistic feedbacks favour body-size stasis but erode genetic variability and associated body-size evolvability, and may ultimately impair population persistence and recovery. In contrast, synergistic feedbacks drive fast evolution towards smaller body sizes and favour population resilience, but may have far-reaching bottom-up or top-down effects. We illustrate the further complexities resulting from multiple environmental feedbacks using a co-evolving predator-prey pair, in which case outcomes from EEFLs depend not only on population densities, but also on whether prey sit above or below the optimal predator/prey body-size ratio, and whether prey are more or less evolvable than their predators. EEFLs improve our ability to understand and predict nature’s response to harvesting, but their integration into the research agenda will require a full consideration of the effects and dynamics of natural selection.

## Introduction

The management of exploited populations is classically based on density-dependent population models in which harvesting, while decreasing population size, also relaxes density-dependent competition so that individual biomass productivity is increased (Verhulst 1838; Schaefer 1954; Hilborn and Walters 1992). However, this classical view has been repeatedly challenged by studies showing that individual biomass productivity often tends to decrease, not to increase, with harvesting (Kristiansen and Svåsand 1998; Conover and Munch 2002; Swain et al. 2007; Edeline et al. 2007, 2009; Biro and Post 2008; Heino et al. 2013).

This negative relationship between harvest effort and individual biomass production is generally interpreted as a rapid evolutionary response to direct harvest selection against large-bodied individuals by fishers (Trippel 1995; Law 2000; Kuparinen and Merilä 2007; Fenberg and Roy 2008; Heino et al. 2015). Accordingly, selection against a large body size is expected to favour slow-growing and early-maturing genotypes, which also tend to have lower fecundity and decreased offspring quality (Walsh et al. 2006; Heino et al. 2013). However, cases remain where exploitation induces no phenotypic change (Hilborn and Minte-Vera 2008; Devine and Heino 2011; Silva et al. 2013; Marty et al. 2014), or a change towards larger body sizes as predicted by density-dependent population models (Hilborn and Minte-Vera 2008). Therefore, whether harvest-induced evolutionary changes occur at all, or are large and rapid enough to influence biomass productivity remains controversial (Andersen and Brander 2009; Borrell 2013).

This debate, we feel, is plagued by a pervasive inclination of many researchers to overlook natural selection and to consider selective removal by harvesters as the only dynamic selective force at play. Natural selection, if ever mentioned, is regarded as negligible such that harvest-induced changes are widely considered as slowly reversible (see e.g. the seminal paper by Law 2000). The simplifying assumption that natural selection is negligible further leads to conclude, for instance, that body-size stasis in harvested populations indicates that evolution is absent or has unimportant effects relative to the effects of ecology (e.g., Hilborn and Minte-Vera 2008). As we propose below, body-size stasis may in fact reflect eco-evolutionary dynamics in which natural selection opposes the effects of direct harvest selection. Failure to account for these eco-evolutionary processes might fundamentally hamper our ability to understand and, hence, to manage productivity dynamics in harvested populations.

The objective of this essay is to provide an impetus to the study of natural selection in harvested populations, and may be seen as a complementary follow-up to Kinnison et al. (2015). These authors stressed that our inclination to seek for parsimonious explanations may drive us to overlook the action of complex eco-evolutionary dynamics. Following this idea, we review theoretical, experimental and empirical insights to elaborate plausible scenarios under which complex eco-evolutionary dynamics may affect the yield and resilience of harvested populations in outcomes that may appear very similar to purely ecological or purely evolutionary dynamics. Most of our examples come from fisheries which, because of their ecological, economic and social importance, concentrate the majority of the literature devoted to harvested animal populations.

We first review the mechanisms through which natural selection may favour either large-bodied or small-bodied individuals. Secondly, we build on this knowledge of natural selection to identify the different pathways and directions that size-dependent EEFLs may take in harvested populations when only one single species evolves. In the third section, we extend the approach to EEFLs acting at the two-species and food-web levels. Finally, in the fourth section we conclude with an overview of the methods currently available to advance our empirical knowledge of EEFLs and with a consideration of how EEFLs may change our approach to managing harvested populations.

### 1. Size-dependent natural selection

Ample evidence shows that trait evolution in response to natural selection may be large and fast, hence far from negligible (Hendry and Kinnison 1999, 2001; Grant and Grant 2002; Stockwell et al. 2003; Hairston et al. 2005; Carroll et al. 2007). This section provides details of the mechanisms through which natural selection moulds body sizes.

#### 1.1 Natural selection for a smaller body size

Competition may be exploitative, i.e., resource-mediated (or indirect) or interference-mediated, i.e., direct. Both types of competition are expected to generate selection on body size, but only exploitative competition is expected to favour smaller body sizes. Exploitative competition may be usefully construed using the *R*\* rule, which states that competition selects individuals surviving on the lowest equilibrium resource level (Tilman 1982). A lower individual *R*\* (i.e., a higher competitive ability) is achieved by increasing resource intake and/or by decreasing basal metabolic requirements. Note, however, that both resource intake and basal metabolic rate generally increase with body size (Peters 1983; Persson et al. 1998; De Roos et al. 2003; Kooijman 2010). Hence, whether individual *R*\* increases or decreases with body size depends on the relative strengths of allometric constraints acting on resource intake and metabolic rate. If resource intake increases faster with body size than metabolic rate, *R*\* decreases with increasing body size and exploitative competition should select for larger body sizes. In contrast, if resource intake increases slower than metabolic rate, *R*\* increases with body size and exploitative competition should select for smaller body sizes. In fish, available evidence suggests that *R*\* increases with body size (Persson and De Roos 2006), so that exploitative competition should favour smaller sizes. The argument extends to many other taxa if one assumes that ingestion increases with body surface (*∝* size^2^) while maintenance increases with body volume (∝ size^3^) (Kooijman 2010). Population dynamics consistent with this prediction have been reported in the vendace *Coregonus albula* (Hamrin and Persson 1986), roach *Rutilus rutilus* (Persson et al. 1998) and Japanese medaka *Oryzias latipes* (Edeline et al. 2016).

Competition, if not leading to competitive exclusion, may also select on body sizes indirectly through decreasing the individual resource share on the long term. Available evidence suggests that such food stress has opposite effects on somatic growth rate and age at maturation. Across a wide variety of taxa, food stress favours slower growth rates and smaller size at age, presumably by imposing energy reallocation to the most vital functions (Arendt 1997). In contrast, fitness-maximising models predict that food stress should select for delayed maturation and, hence, for larger size at maturity if somatic growth rate is constant (Gadgil and Bossert 1970), a prediction supported by available empirical evidence (Holliday 1989; Sgrò and Partridge 2000). Therefore, if somatic growth and maturation trade off, the growth-mediated and maturation-mediated effects of food stress on body size oppose each other, and are thus likely to remain inconsistent or cryptic.

Competition is not the only ecological interaction that may select for smaller body sizes. Predators that target large-bodied prey directly select for smaller prey body sizes just like harvesters do (see above). This is for instance the case for fish predation on zooplankton (Brooks and Dodson 1965). If predators are non size-selective, predators still favour earlier maturation in prey and, hence, a smaller size at maturity if somatic growth is constant (Abrams and Rowe 1996). This is because early-maturing individuals have an increased fitness advantage when life expectancy is reduced (Gårdmark and Dieckmann 2006; Heino et al. 2015). Finally, if predation mortality is stage-dependent, higher juvenile (immature) mortality favours earlier maturity which, given a fixed somatic growth rate, also means maturity at a smaller body size (Abrams and Rowe 1996; Heino et al. 2015).

#### 1.2. Natural selection for a larger body size

In the wild, survival often increases with larger body sizes (Roff 1992), as demonstrated for instance in juvenile fish (Perez and Munch 2010; Stige et al. 2019), juvenile Soay sheep (*Ovis aries*, Hunter et al. 2018), or adult fish (e.g., Carlson et al. 2007, Olsen and Moland 2011). The mechanism behind this positive survival-size relationship could involve a higher resistance to starvation in larger-bodied individuals (van de Wolfshaar et al. 2008), but also results from strong interference in competitive interactions. While size-selective effects of exploitative competition are dependent upon the allometric scaling exponents of intake and maintenance rates (see above), interference competition almost universally brings an advantage to large-sized individuals in contests for food (Persson 1985; Post et al. 1999). In fish, dominance hierarchies are highly size-dependent both among and within species (Griffiths et al. 2020; Fausch et al. 2021). In experimental populations of the springtail *Folsomia Candida*, interference favours large-sized individuals that can monopolize resources (Le Bourlot et al. 2014). Similarly, in wild populations of the brown anole lizard *Anolis sagrei* natural selection for larger body sizes increases in parallel with population density and associated interference competition (Calsbeek and Smith 2007). In these lizards, the strength of competition-induced selection on body size overwhelmed the strength of predation-induced selection (Calsbeek and Cox 2010).

Often, predators are size-limited and thus preferentially feed on small-sized prey individuals. This is true for both aquatic and terrestrial systems (Sinclair et al. 2003). In such cases, predators favour prey individuals that grow fast through a “predation window” to rapidly reach a size refuge, i.e., they select for a large body size at a given age (Day et al. 2002). This predation window plays a key role in mediating the population dynamic effect of intraspecific predation (i.e., cannibalism), an interaction that is present in multiple aquatic or terrestrial taxa (Fox 1975; Claessen et al. 2002, 2004). Cannibalism is presumably the mechanism that controlled the positive effect of population density on somatic growth rate in the pike (*Esox lucius*) population of Windermere where, as the density of cannibals increased, survival was biased towards faster-growing individuals (Edeline et al. 2007, 2009).

The effect of size-limited predation on age at maturation is less straightforward than on somatic growth. If mortality increases among small-sized individuals, predictions depend on the details of the model. Optimality models predict evolution of delayed maturation at a larger body size (Taylor and Gabriel 1992). In contrast, adaptive dynamics models accounting for a trade off between somatic growth and reproduction and for a positive effect of body size on fecundity lead to more complex outcomes: increased mortality among small-sized individuals can increase or decrease maturation size, or even lead to the coexistence of both early- and late-maturing individuals when benefits from early maturation collide with benefits from growing fast to a size refuge (Gårdmark and Dieckmann 2006). The outcome of evolution also depends on whether food availability is sufficient to support fast somatic growth (Chase 1999). To our knowledge, the available empirical and experimental evidence more often supports delayed maturation at a larger body size when predation targets small-sized individuals (Edley and Law 1988; Wellborn 1994; Beckerman et al. 2010; Le Rouzic et al. 2020).

Finally, a larger body size further provides females with a higher fecundity in egg-spawning species (Barneche et al. 2018), and males with a strong advantage in contest sexual selection (e.g., Fleming and Gross 1994). Combined together, these multiple positive effects of natural selection on body sizes are likely to outweigh the negative effects, as suggested by an overall tendency for natural and sexual selection to favour larger body sizes across multiple taxa (Kingsolver and Pfennig 2004). We now move to examining how such size-dependent natural selection may interact dynamically with ecology in EEFLs.

### 2. Theory and scenarios of harvest-induced EEFLs with one evolving species

#### Box1. Defining the selection- and evolvability-mediated pathways to eco-evolutionary feedback loops (EFFLs).

To study existing feedbacks between ecological and evolutionary dynamics, two main frameworks are currently used: quantitative genetics (QG) and adaptive dynamics (AD). Though the two methods differ, they are both based on the idea that the description of trait dynamics in response to selection requires two fundamental ingredients: trait(s) evolutionary potential (hereafter “evolvability”) and a measure of selection cting on the trait(s) (Abrams 2001). Consider for instance the classical recursive equation of quantitative genetics (QG):

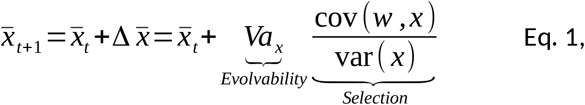

where 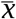 is the mean population value of a univariate trait *x, t* is generation index, *Va_x_* is additive genetic variance, *w* is relative individual fitness, and 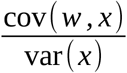 is the directional selection gradient, i.e., the slope of the linear regression between relative fitness and trait *x* (Lande and Arnold 1983). Provided that the definition of *w* includes at least density dependence and/or frequency dependence, Eq. 1 incorporates selection-mediated EEFLs as the ecological context (density or frequency) then impacts the selection term (Abrams 2001). Eco-evolutionary feedback loops may also occur through the evolvability-mediated pathway in Eq. 1, for instance if *Va_x_* is directly linked to the demographic context (e.g., an existing correlation between population density and genetic variability) or if *Va_x_* is an explicit function of the strength of selection since strong directional selection is expected to decrease additive genetic variances (Crow 2008).

Adaptive dynamics (AD) (Dieckmann and Law 1996) readily account for both selection- and evolvability-mediated EEFLs. This essential feature of AD is captured by the canonical equation:

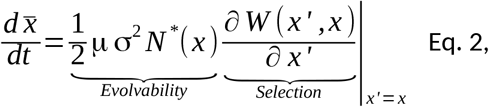

where *x* is a resident trait, *x’* is a mutant trait, 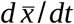 is a continuous-time analogue of 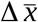 in Eq. 1, μ is per capita mutation rate, and σ^2^ is phenotypic variance from a mutation. *N*\* (*x*) is equilibrium population size for the resident trait, and Eq. 2 hence incorporates the evolvability-mediated pathway to EEFLs since evolvability, here determined by the mutation process, is explicitly dependent on equilibrium population size *N*\* (*x*), which is set by the value of the resident trait *x. W*(*x′,x*) is invasion fitness for a mutant trait *x′* in an environment determined by the resident trait *x*. Because this fitness definition is based on ecological dynamics, one sees that selection-mediated EEFLs are readily considered in adaptive dynamics models. Finally (*∂ W* (*x′,x*))/(*∂ x′*) is the directional selection gradient acting on the mutant trait *x′*, i.e., is the invasion criterion (slope of the fitness landscape for *x′* evaluated in *x*).

Theory presented in Box 1 predicts that EEFLs may proceed through two different pathways: a selection-mediated and an evolvability-mediated pathways, which we illustrate in Fig. 1. The selection-mediated pathway is captured by Arrow 1 (Fig. 1): the environment of an individual generates natural selection on body size (see Section 1 above). In addition to influencing individual fitness and, from there, population densities, body size has widespread and consistent ecological effects (Peters 1983; Brown et al. 2004; Woodward et al. 2005). Hence, selection-induced change in body size, in turn, may impact the environment through the size-dependency of reproductive success and ecological interactions (Arrow 2). The evolvability-mediated pathway to size-dependent EEFLs is captured by Arrow 3, and involves mutation-limitation effects linked to population sizes and/or any other existing correlations between genetic diversity and population size (Box 1, Frankham 1996). Such evolvability-mediated pathways to EEFLs are often neglected, but may actually be important from a management or a conservation point of view (Carlson et al. 2014; Marty et al. 2015; Kuparinen and Hutchings 2017). Harvesting may trigger or disrupt size-dependent EEFLs through both direct harvest selection on body size and through the removal of conspecifics and possibly also heterospecifics (Fig. 1).

**Figure 1:**
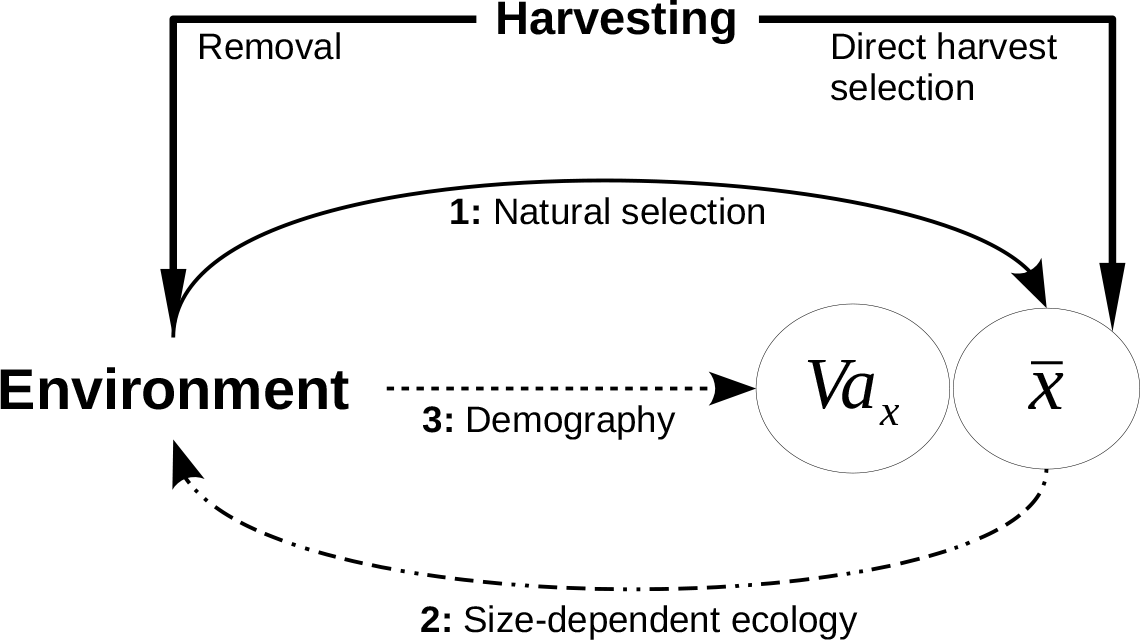
Selection-mediated and evolvability-mediated pathways to size-dependent eco-evolutionary feedback loops (EEFLs). *Va_x_* and 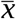 stand for additive genetic variance and mean body size, respectively, in the harvested population (Box 1). Arrow 1: natural I selection and the selection-mediated pathway to EEFLs, Arrow 2: body size-dependent ecological processes, Arrow 3: Effects of demography on genetic variability and the evolvability-mediated pathway to EEFLs (Box 1). NB 1: phenotypic plasticity, as also captured by Arrow 1, will not be discussed. NB 2: For the sake of simplicity, we did not consider the potential direct effects of selection on additive genetic variances (e.g. Crow 2008).

In order to fully grasp the basic ideas that underpin size-dependent EEFLs in the system depicted by Fig. 1, we provide a graphical representation of a moving adaptive landscape in Fig. 2. For simplicity, the fitness landscapes represented on fig 2 ignore frequency dependent selection, so that each phenotype has a given fitness irrespective of its frequency. This fitness would be representative of the absolute fitness of the corresponding monomorphic population. Relative fitness of this phenotype confronted to another can then simply be read from the relative position on the fitness landscapes, phenotypes with higher fitness being selected. Therefore this representation, though it simplifies the underlying ecological aspects, allows to assess absolute and relative fitness easily. We make this choice because population persistence depends on absolute, not relative fitnesses, and because absolute fitness is therefore more intuitively linked with management aspects. For more discussion on the link between absolute and relative fitness, see Orr (2007).

**Figure 2:**
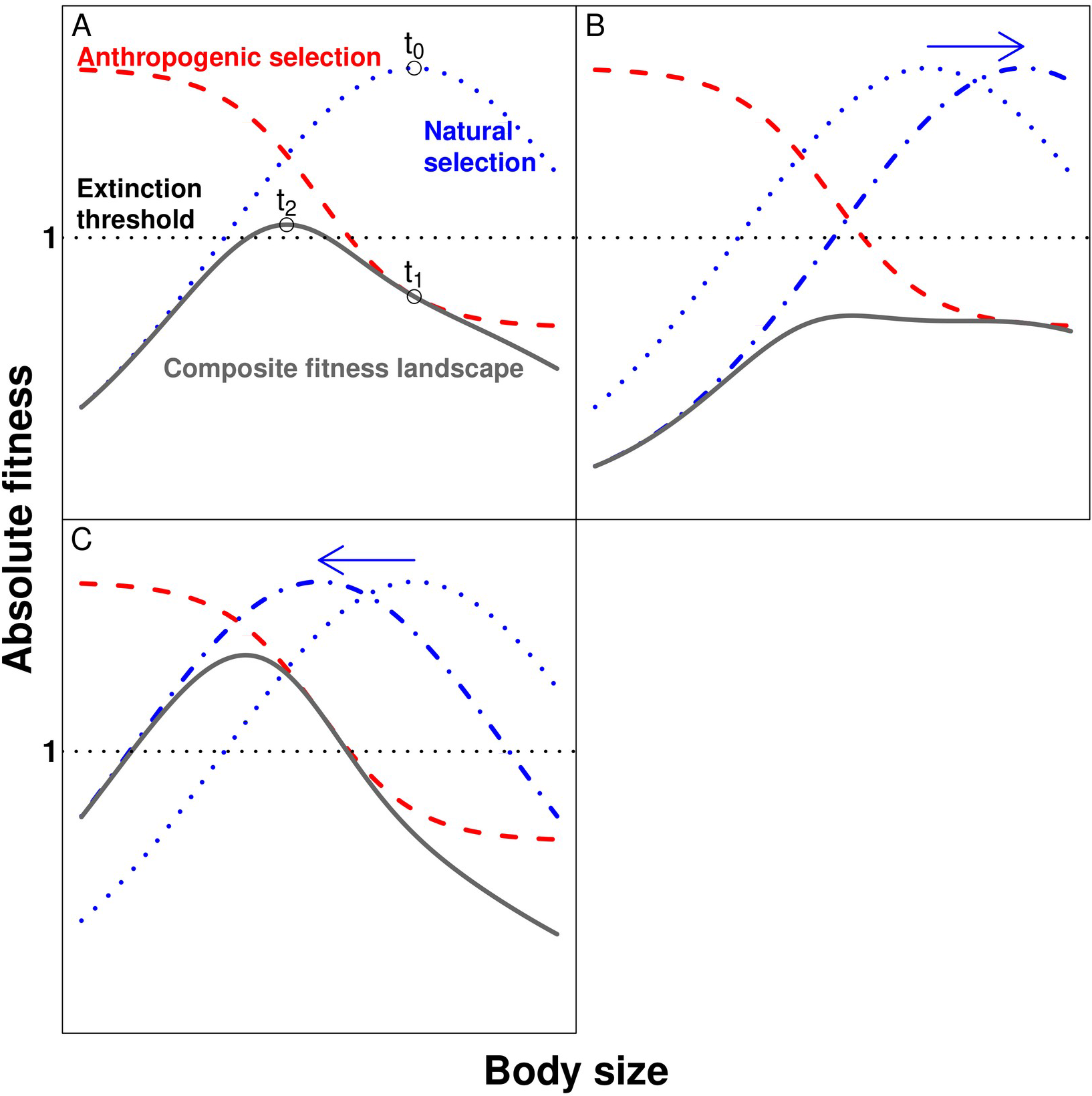
Eco-evolutionary feedbacks in harvested populations. Curves show the relationship between absolute fitness and body size, and the horizontal dotted line shows unity absolute fitness (extinction threshold). **A:** Evolutionary “rescue” (see Glossary) with no eco-evolutionary feedback loop (EEFL). Open circles show the mean phenotype in the population. **B:** A single density parameter feedbacks on natural selection, generating an antagonistic EEFL. The arrow shows the change in directional natural selection due to the environmental feedback. **C:** The one-dimensional density-dependent feedback generates a synergistic EEFL in which natural selection changes to reinforce the effect of direct harvest selection.

Second, we represent what we believe are the most generic functional forms for absolute fitness functions: harvest selection is often directional against a large body size, and body size is often constrained by trade-offs resulting in stabilizing natural selection (Kingsolver et al. 2001, Kingsolver and Pfennig 2004, Carlson et al. 2007). Note that this assumption of stabilizing natural selection also follows from the general observation that evolution towards smaller body sizes is associated with severe fitness costs (e.g., Walsh et al. 2006), while largest-bodied and oldest individuals may be subject to senescence. Although these settings are very simple and maybe rare in nature, their evolutionary outcome is more easily visualized than when multiple environmental feedbacks operate simultaneously and prevent evolutionary optimization (Metz et al. 2008).

In the absence of any direct harvest selection, the population mean body size resides at the naturally-selected body size optimum (dotted blue curve, t_0_ in Fig. 2A). The product of natural selection with direct harvest selection (i.e., survival to harvesting, dashed red curve) instantaneously warps the naturally-selected fitness landscape to generate a new, composite fitness landscape (solid grey curve) on which the population mean trait value is associated with a fitness at which the population crosses the extinction threshold (t_1_, maladaptation). Rapid adaptive evolution through a few generations towards the newly-selected adaptive optimum restores a fitness at which the population may persist (t_2_, *re*-adaptation). If adaptive change occurs fast enough, it may potentially restore a positive population growth and prevent extinction, a process termed “evolutionary rescue” (Glossary, Gomulkiewicz and Holt 1995).

The model presented in Fig. 2A makes the simplifying assumption that natural selection does not respond to harvesting. However, in addition to imposing direct harvest selection on body size, harvesting also alters the environment (Fig. 1, Arrow 1) and may thus indirectly change natural selection acting on body size (Bouffet-Halle et al. 2021). We will now examine two scenarios in which such harvest-induced changes in natural selection either oppose or reinforce the action of direct harvest selection on body size.

#### 2.1. Antagonistic EEFLs

We first consider a feedback in which harvesting changes natural selection towards favouring *larger-than-initial* body sizes (sketched in Fig. 2B). As this selection acts in opposite ways to the direct selective effects of harvesting, we refer to this situation as an **antagonistic EEFL**. Compared to an EEFL-absent case, antagonistic EFFLs magnify warping of the adaptive landscape and thus impair population persistence. At an extreme, the fitness peak may dwindle below the extinction threshold (Fig. 2B). Antagonistic EEFLs are expected whenever density-dependent natural selection favours small body sizes and harvesting, through reducing densities, relaxes natural selection for a small body size (Fig. 2B). For instance, reduced population densities may relax exploitative competition for resources, and weaken associated selection for smaller body sizes (Table 1). Antagonistic EEFLs may also emerge from changes in predation regimes, as demonstrated by Gårdmark et al. (2003) using a theoretical model in which an age-structured population evolves in response to both harvesting and predation mortality. Harvesting reduces prey availability so that predator density decreases, thus inducing relaxed predation and the associated natural selection for smaller body sizes. This result is likely to apply whenever predators of the harvested population directly select for smaller body sizes, i.e., when predators preferentially prey on large-bodied individuals, on juveniles, or when they are non size-selective (Table 1).

**Table 1.**
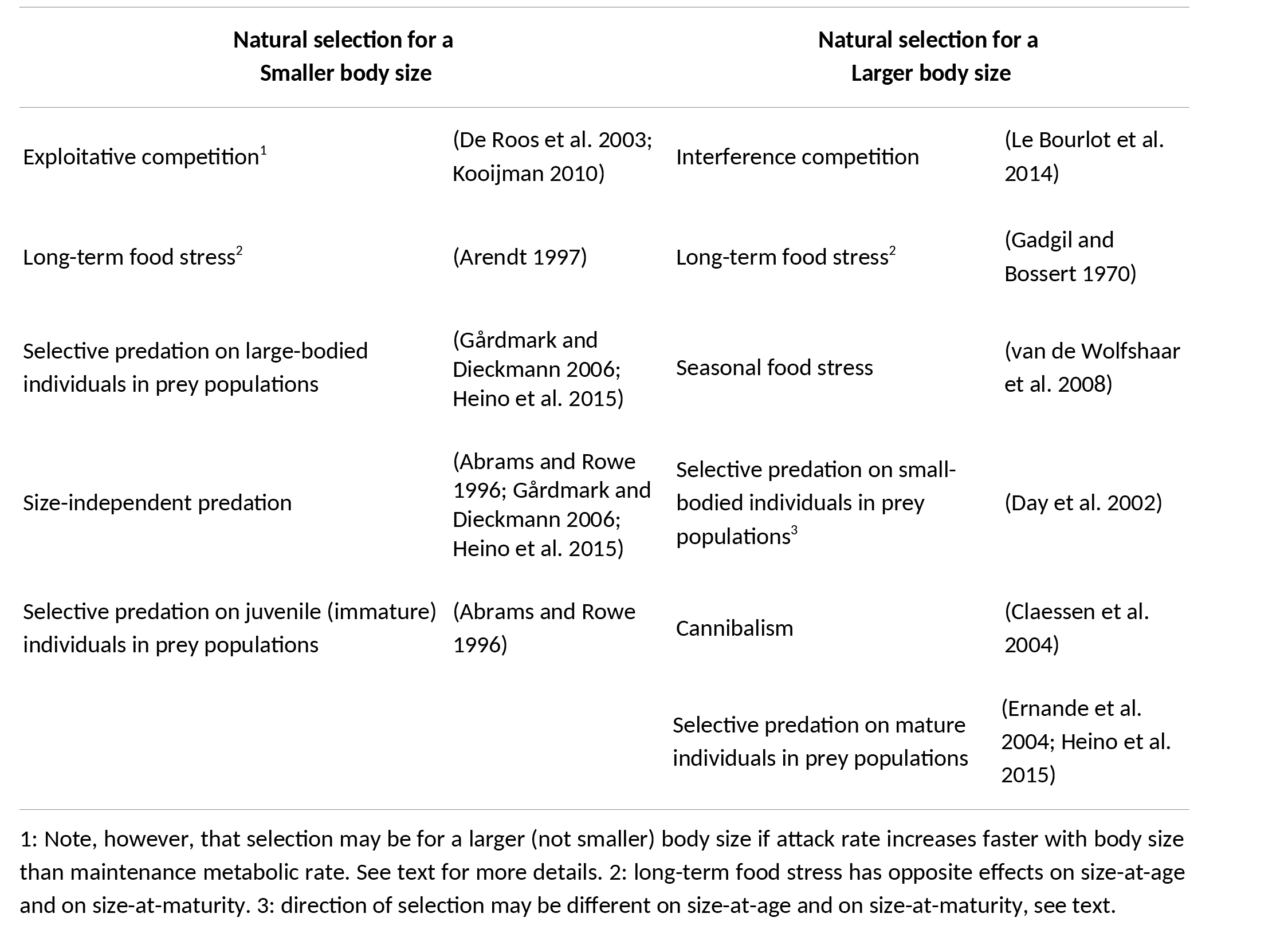
Sources of natural selection predicted and observed to favour either a smaller or larger body size at age or at maturity.

A hallmark of eco-evolutionary dynamics is their tendency to remain cryptic if they are not anticipated and, hence, not specifically investigated (Kinnison et al. 2015). Size-dependent, antagonistic EEFLs are no exception, because the changes in natural selection oppose the effects of direct harvest selection and favour body-size stasis, an outcome that may erroneously be interpreted as direct harvest selection being too weak to drive any evolutionary response (e.g., Hilborn and Minte-Vera 2008). In fact, however, body-size stasis of antagonistic EEFLs is associated with a fitness drop that may ultimately prevent evolutionary rescue (Fig. 2B). The fitness drop and resultant decreased population size may further jeopardize body-size evolvability (Box 1, Arrow 2 → 3 sequence in Fig. 1) which, together with a vanishing strength of selection due to a flat composite fitness landscape (Fig. 2B), decreases the probability for recovery. Overall, any situation in which harvesting is associated with body-size stasis but severe population decline may be suspected to reflect an antagonistic EEFL.

#### 2.2. Synergistic EEFLs

**Synergistic EEFLs** occur when the environmental feedback changes natural selection towards favouring smaller-than-initial body sizes in synergy with direct harvest selection (Fig. 2C). Synergistic EEFLs may result, for instance, when harvesting, through reducing the density of large-sized individuals in the population, relaxes interference competition and cannibalism and associated natural selection for a large body size (Table 1). Recent experimental evidence in replicated fish populations suggests that this harvest-induced relaxation of interference competition and cannibalism can drive a rapid evolutionary divergence between harvested and non-harvested populations (Bouffet-Halle et al. 2021). Synergistic EEFLs are also expected when predation favours larger body sizes and predators disappear due to harvest-induced prey shortage (Table 1, Jusufovski and Kuparinen 2020).

Qualitatively, the phenotypic outcome from synergistic EEFLs looks similar to the phenotypic outcome from EEFL-absent dynamics (Fig. 2A), though directional selection is stronger and expected trait variation faster. Synergistic EEFLs are thus likely to remain cryptic and to be interpreted as a large and rapid response to direct harvest-selection acting alone (e.g., Darimont et al. 2009). Compared to an EEFL-absent situation (Fig. 2A), however, synergistic EEFLs result in a magnified fitness peak on the composite fitness landscape (Fig. 2B) and, hence, favour larger population sizes at the body-size optimum and higher body-size evolvability (Arrow 2 → 3 sequence in Fig. 1). Hence, synergistic EEFLs may favour evolutionary rescue and allow fast evolutionary rebound after relaxation of fishing. This is presumably the configuration that explains why pike, a highly cannibalistic species, showed a fast and large evolutionary response to varying harvesting intensity in Windermere (Edeline et al. 2007, Coltman 2008). Finally, synergistic EEFLs increase slope steepness around the fitness peak on the composite fitness landscape (Fig. 2C), resulting in stronger selection around and faster evolution towards the body-size optimum. Therefore, synergistic EEFLs are consistent with the observation that fishing-induced trait changes are often much faster than predicted by theoretical models that only assume direct harvest selection (Audzijonyte et al. 2013a).

These simple scenarios of antagonistic and synergistic EEFLs focus on the evolution of just one harvested species alone. Fisheries, however, most often target not just one but several species within the ecological network, so that an ecosystem perspective on fishery management is required (White et al. 2012; Perälä and Kuparinen 2020). Therefore, we now move to examining EEFLs when more than one species evolves.

### 3. Scenarios of harvest-induced EEFLs with multiple evolving species

There is currently an emerging recognition that evolution in a given harvested species can induce coevolution in other species through changes in ecological interactions (Wood et al. 2018). However, understanding the evolutionary response to harvesting in a multispecific context is highly challenging (Audzijonyte et al. 2013b). Investigation on multispecies EEFLs requires to account simultaneously for the coevolution of the various body sizes, of the network structure, and to consider how one feeds back on the other (Loeuille and Loreau 2005). Direct data investigating the occurrence and magnitude of multispecies EEFLs are scarce. However, different empirical facts suggest that multispecies EEFLs may naturally emerge in exploited ecological networks.

First, empirical data suggest that predators are often larger than their prey in both terrestrial and aquatic systems (Cohen et al. 2003; Sinclair et al. 2003; Brose et al. 2006) and that predator-prey body-size ratios determine the strength of predation (Emmerson and Raffaelli 2004; Renneville et al. 2016). Therefore, we expect that selection on body size will change the distribution of interaction strengths, which largely constrains ecosystem functioning and stability (e.g., McCann et al. 1998, Rooney et al. 2006). Ultimately, rewiring and redistribution of interaction strengths may lead to extinctions in the network. For instance, evolution of larger body sizes can decrease the density of the evolving population thereby increasing its vulnerability to demographic stochasticity and potentially facilitating its extinction (evolutionary deterioration, see Glossary). At the same time, body-size evolution may undermine predator persistence through weakening trophic links, or compromise prey persistence through strengthening trophic links. Similarly, variations in interaction strengths will affect apparent competition (Holt et al. 1994), thereby changing coexistence conditions within the food web and possibly leading to competitive exclusions. Because the network structure, in turn, constrains the fitness of species within the community, multispecies EEFLs naturally emerge (Fig. 3).

**Figure 3:**
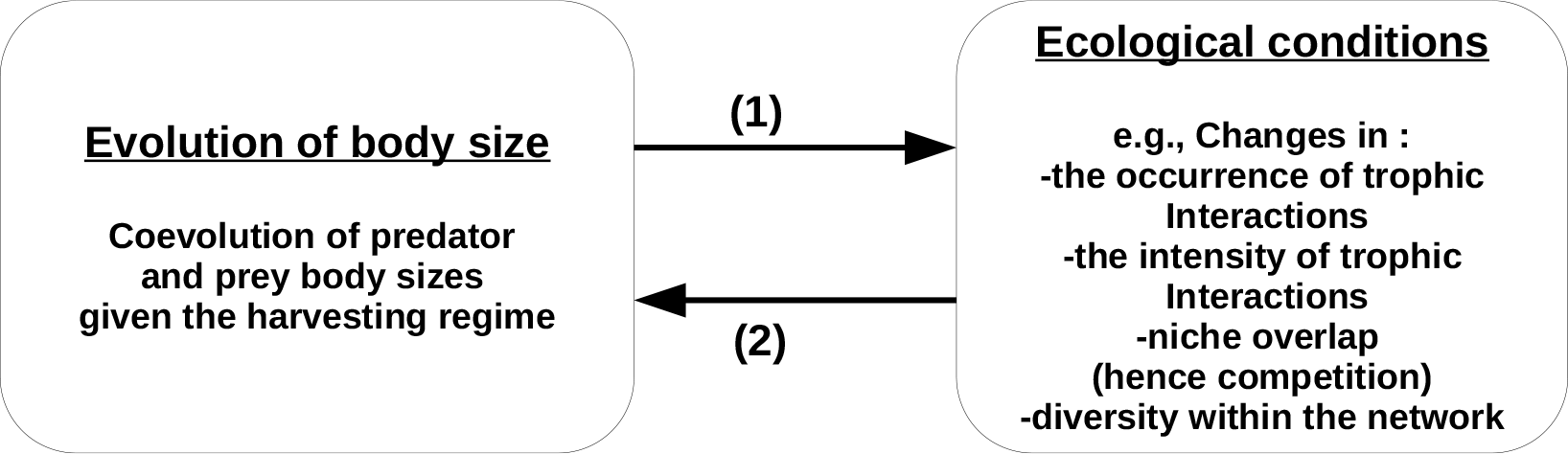
Components of multispecies EEFLs in food webs. (1) Coevolution of body sizes within the network under the new selective regime (harvesting) affects interaction patterns within the network, as well as coexistence conditions. (2) This new ecological context in turn changes the fitness of species (eg, due to changes in predation rates), thereby acting on body size coevolution.

Co-evolution strongly complicates EEFLs. In particular, the graphical framework from Fig. 2 no longer applies, because the environment now becomes multidimensional and evolution no longer optimizes fitness or population size of any given species (Meszéna et al. 2001; Metz et al. 2008). To keep our arguments as simple as possible we focus on a single co-evolving predator-prey pair, in which we examine two non-exclusive mechanisms for the emergence of EEFLs. We first examine the consequences of a “trophic relaxation”, which occurs when decreased densities weaken the strength of the predator-prey link. Second, we examine the consequences of predators and prey having “asymmetric evolvability” for body size and, hence, evolving at a different pace in response to harvesting. For both mechanisms we consider that, before harvesting starts, the predator-prey pair resides at an evolutionary equilibrium.

#### 3.1. Trophic relaxation

Prey may be either smaller or larger than the preferred prey size of the predator. Prey sizes matching the preferred size are rarely expected, as prey may evolve away from such situations, but also because the distribution of body sizes does not usually maximize trophic interactions due to metabolic constraints, competition and the multiplicity of prey and predator species that also act as selective pressures (e.g., Loeuille and Loreau 2005, 2006).

We now start harvesting both the prey and predator which, hence, are both under direct harvest selection for a smaller body size (red arrows in Fig. 4). In Figs. 4A and 4B, red arrows have similar lengths indicating that both species evolve smaller body sizes at a similar pace, such that no change is to be expected in their realized body-size ratio. However, because harvesting reduces population density in both the prey and predator, we expect a relaxation in the strength of the predator-prey link. This “trophic relaxation” is outlined in Figs. 4A and 4B by a decreased predation intensity (dotted Gaussian curves). Such a trophic relaxation may lead to opposite eco-evolutionary outcomes depending on whether initial prey size is smaller or larger than the predator’s optimal prey size.

**Figure 4.**
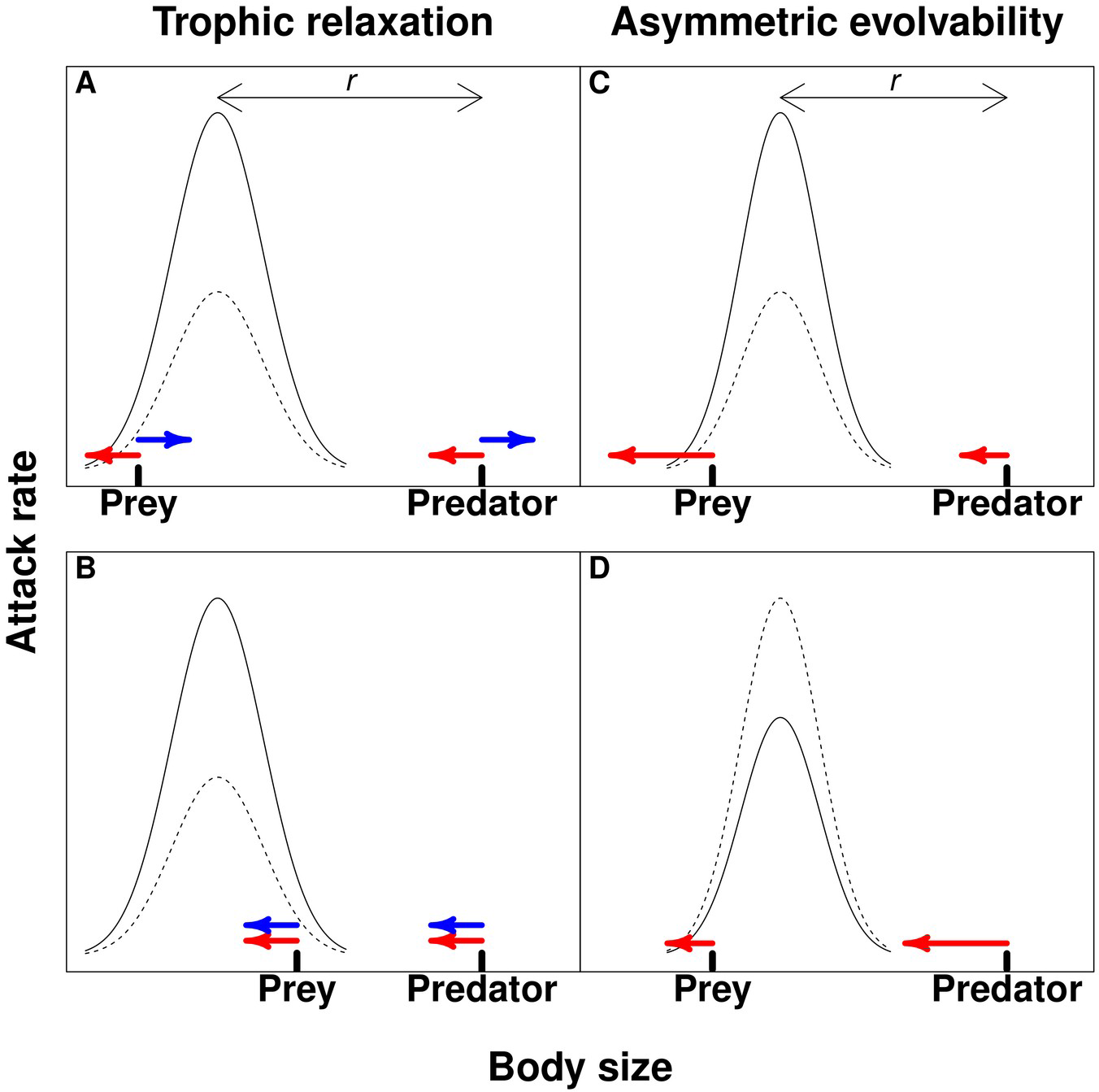
Alternative settings in coevolving predator-prey pairs. The Gaussian curves show the predation intensity before (solid lines) and after (dashed lines) harvesting starts. Optimal prey size is set by the fixed ratio *r*. Red horizontal arrows show potential for body size evolution (i.e., body-size evolvability) in response to direct harvest selection. Blue arrows show natural-selection response to harvesting, i.e., the EEFL. **A**: Trophic relaxation leading to increased natural selection for a larger body size. **B**: Trophic relaxation leading to natural selection for a smaller body size. **C**: Body-size evolvability is larger in the prey than in the predator. **D**: Body-size evolvability is larger in the predator than in the prey.

In Fig. 4A, prey size is initially smaller than optimal for the predator, and the trophic relaxation thus results in relaxed natural selection for smaller body sizes in both the prey and predator (blue arrows). In other words, change in natural selection acts in opposition with direct harvest selection in an antagonistic EEFL. In case 4B, in contrast, prey size is initially larger than optimal predator size, and the trophic relaxation thus results in relaxed natural selection for larger body sizes, thus creating a synergistic EEFL. Of course, these relatively simple outcomes are complicated by feedbacks from intraspecific interactions (competition, cannibalism) that may either reinforce or oppose the effects of the predator-prey feedback (see Section 2).

#### 3.2. Asymmetric evolvability

Assuming symmetric evolvability in the prey and predator (Figs. 4A and 4B) is likely unrealistic for most situations. Rather, body-size may be more evolvable in prey than predators (hence the longer red arrow in Fig. 4C), either because the trait is determined by different gene networks for the two species, or because the two species have very different population sizes, hence differing in accumulation of mutations or standing genetic variability. For instance, smaller (prey) body sizes are often associated with larger population numbers (Woodward et al. 2005) and with a higher genetic variability (Romiguier et al. 2014; De Kort et al. 2021). Under these settings, prey evolve smaller body sizes faster than their predator, move further away from predator’s preferred prey size, and ultimately benefit from an evolution-induced trophic relaxation (Fig. 4C). The predator on the other hand, may become resource limited, so that further declines in predator population are expected. This is different from trophic relaxation in cases 4A and 4B which was the driver of evolution.

In Fig 4D, we sketch an opposite, perhaps less common situation in which predators have a higher body-size evolvability than their prey. This configuration may potentially result from prey being close to a lower evolutionary limit for body size (Le Rouzic et al. 2020; Renneville et al. 2020). Under these settings, predators evolve smaller body sizes faster than prey, such that preferred prey size moves closer to prey size and a trophic magnification results. Such a coevolution therefore favours the maintenance of the trophic interaction. Note that these outcomes depend on prey being smaller than predator’s preferred prey size in Figs. 4C and 4D, and are reversed when prey are larger than the preferred prey size of the predator (i.e., trophic magnification in Fig. 4C and trophic relaxation Fig. 4D).

#### 3.3. More complex interaction networks

In more complex networks, the multiplicity of trophic and non-trophic interactions may generate a variety of counteracting selection gradients, so that evolution might be more constrained than in a single predator-prey link. If this hypothesis is true, EEFLs might well be more important in explaining evolutionary and ecological stasis rather than change (Ellner et al. 2011; Strauss 2014; Kinnison et al. 2015). Beyond very specific scenarios, network and eco-evolutionary complexities under harvesting scenarios are virtually impossible to grasp intuitively, and are even hard to handle through a mathematical analysis. However, numerical simulations are certainly possible. In this regard, the development of evolutionary models of food webs based on body size offer promising venues, as they already consider simultaneously evolution of body size and changes in the network structure (Loeuille and Loreau 2005, 2009; Brännström et al. 2011; Allhoff et al. 2015). Harvesting scenarios may be implemented in such models (Perälä and Kuparinen 2020), as has been done in other contexts (eg, climate warming, Weinbach et al. 2017, Yacine et al. 2020).

### 4. Management perspectives

So far, the vast majority of models used to project the eco-evolutionary consequences of fishing ignore natural selection on body size (but see Jusufovski and Kuparinen 2020). Hence, although quite elaborated, these models are likely to underestimate either the demographic consequences of harvesting when antagonistic EEFLs are involved, or the rates of evolutionary change and recovery when synergistic EEFLs are involved. We recognize, however, that more empirical and experimental studies are needed to document the pathways, directions and strength of density-dependent selection acting on body size in harvested systems. In particular, it is important to document whether and when harvest-induced EEFLs can be simplified into a one-dimensional, density-dependent process that can be handled by optimality approaches such as that outlined in Fig. 2. In Box 2, we provide an overview of the empirical methods currently available to progress in that direction.

#### Box 2. Empirical exploration of size-dependent EEFLs: where to go next?

Demonstrating a full selection-mediated EEFL requires showing both that natural selection drives evolutionary trait change and, in turn, that the resultant trait evolution alters the environment in such a way that natural selection acting back on the trait is modified (Figs. 1, 2). Considering also the evolvability-mediated pathways to EEFLs requires to further measure the effects of environmental changes on trait evolvability. Tackling such a complexity is challenging but, as we show below, not beyond of reach.

##### Measuring natural selection and trait response to selection

The form and strength of selection are most accurately measured by estimating fitness-traits relationships at the individual level (Arnold 2003) using, e.g., capture-recapture techniques. Alternatively, the directional component of selection may also be estimated from population and trait time series using the “Geber method”, the age-structured price equation or integral projection models (Hairston et al. 2005; Ellner et al. 2011; van Benthem et al. 2017; Govaert 2018). A drawback of all these methods is that they measure selection acting on phenotypes, while evolution is concerned only by selection acting on the heritable component of phenotypes (Morrissey et al. 2010). To circumvent this problem, statistical approaches making use of the “animal model” (AM) of quantitative genetics were developed to specifically measure selection acting on the additive genetic component of traits and, hence, to accurately predict evolution (Hadfield 2008; Morrissey et al. 2010; Stinchcombe et al. 2014). AM-based approaches require pedigree data and are thus more readily implementable in small, closed systems than in large-scale fisheries (but see Koch et al. 2008).

##### Measuring the dependency of natural selection on the environment (Fig. 1, Arrow 1)

A pivotal condition for the emergence of selection-mediated EEFLs is that natural selection dynamically changes due to changes in the environment (Figs. 1, 2, Govaert et al. 2019). This may be checked *a posteriori* through measuring genotype-by-food interactions on body sizes. For instance, Bouffet-Halle et al. (2020) used this approach to show that harvest-induced evolution towards smaller body sizes in experimental populations of medaka fish (*Oryzias latipes*) had evolved in a low-food but not in a high-food environment. This result suggested that medaka had evolved in response to density-dependent natural selection at high population density (low food), but not in response to direct harvest selection at low population density (high food). This approach, however, remains fragile because our understanding of genotype-by-food interactions remains limited, and other complementary results may be necessary to back-up conclusions from genotype-by-food analyses (Bouffet-Halle et al. 2021). When possible, selection-environment relationships should be measured directly using individual capture-recapture techniques (e.g., Haugen et al. 2007, Calsbeek and Smith 2007, Calsbeek and Cox 2010), keeping in mind the problems highlighted above of measuring selection at the phenotype level. Here also, these problems may be solved if the data permits applying the AM, which may be extended to estimate environment-selection relationships acting at the additive genetic level (Hunter et al. 2018).

##### Measuring the trait dependency of ecological dynamics (Fig. 1, Arrow 2)

Time series data may be used to quantify the feedback from phenotypic trait change to environmental variables. Since the inception of the Geber Method by Hairston et al. (2005) and Ellner et al. (2011), a multiplicity of more sophisticated methods have flourished. These methods are based either on inferring parameters for dynamic models from data (e.g., Rudy et al. 2017 and references therein), on non-parametric approaches such as Recurrent Neural Networks, or on hybrid approaches combining differential equations with neural networks (e.g., Bonnaffé et al. 2020 and references therein). Reviewing these methods is beyond the scope of this paper.

##### Measuring the effects of the environment on trait evolvability (Fig. 1, Arrow 3)

Evolvability may be measured using multiple metrics (e.g., Hansen et al. 2011, 2019), which condition approaches to exploring evolvability-mediated EEFLs. Here, we focused on additive genetic variance V_A_ which is a commonly-used measure of evolvability (Box 1), and which we assumed to be positively linked to population size (see Reed and Frankham 2001 for a contrasted view). Estimation of V_A_ relies on the AM, using either pedigrees or genetic markers of coancestry to construct relatedness matrices, with some caveats stressed by Lynch and Walsh (2018). The AM may further be extended to incorporate effects of environmental covariates on V_A_ in so-called random regression approaches (Schaeffer 2004).

Importantly, our review suggests that the ecological and evolutionary consequences of harvesting will largely depend on the ecological factors that regulate the population and, hence, will likely be constrained by the details of the local network context. However, based on our above analysis we may still propose some general management rules accounting for size-dependent EEFLs. As highlighted by Engen et al. (2014), a very general consequence of density-dependent selection is that the more ecologically-sustainable strategies will also produce the less evolutionary changes. Therefore, preventing population declines and alleviating evolutionary change are not independent lines of management but are instead highly intertwined management targets. If possible, management rules should further account for the probability of EEFLs to be either antagonistic or synergistic, because the former are far more detrimental than the later to population persistence and recovery and, hence, would impose lower exploitation rates. Ideally, an *a priori* knowledge of the direction of density-dependent natural selection acting on body sizes could be gained using *had hoc* approaches (Box 2). Alternatively, a basic knowledge of the dominant ecological interactions could be used (Table 1).

In co-evolving predator-prey pairs, managers may also account for body-size ratios and potential asymmetries in body-size evolvability, so as to classify their harvested system into one of the four categories depicted in Fig. 4. Body-size ratios are well documented in the literature, and identification of a context prone to trophic relaxation or magnification should be relatively simple and lead to prudent exploitation. Prudent exploitation is also recommended if asymmetric body-size evolvability is suspected, especially when prey can escape predation (Fig. 4D), a situation in which exploitation rates should be stronger on the faster-evolving species so as to resorb asymmetry in evolvability. This recommendation somehow converges towards “balanced harvesting”, a management approach based on spreading fishing mortality across the widest possible range of species and sizes in proportion to their natural productivity. Interestingly, such balanced strategies have already been advocated to conciliate yield and sustainability even in models that ignore evolution (Tromeur and Loeuille 2017). Although more research is clearly needed to test whether and under which conditions these general recommendations hold true, we believe that far enough evidence is already available showing that a consideration of natural selection is highly needed if we are to improve our ability to accurately predict and manage the dynamics of harvested populations.

## Glossary

Absolute fitness: number of offspring reaching the reproductive stage.
Evolutionary deterioration: evolutionary change leading to smaller population densities, thereby increasing its probability of extinction (e.g., due to demographic stochasticity).
Evolutionary rescue: adaptive evolutionary change that restores positive growth to declining populations and prevents extinction.
Evolvability: trait potential to evolve.
Fitness landscape: multidimensional surface depicting fitness as a function of phenotypic traits.
Relative fitness: absolute fitness of a given phenotype divided by average absolute fitness of all phenotypes in the population.
Selection gradient: Trait-specific slope of the fitness landscape, i.e., holding other traits constant.

## Acknowledgements

EE acknowledges financial support from the Norwegian Research Council (projects EvoSize RCN 251307/F20 and REEF RCN 255601/E40) and from Rennes Métropole (project AIS 18C0356). *Version 4 of this preprint has been peer-reviewed and recommended by Peer Community In Ecology (https://doi.org/10.24072/pci.ecology.100071)*

